# Multiscale Hybrid Fabrication: Volumetric Printing Meets Two-Photon Ablation

**DOI:** 10.1101/2022.10.28.513651

**Authors:** Riccardo Rizzo, Dominic Ruetsche, Hao Liu, Parth Chansoria, Anny Wang, Amelia Hasenauer, Marcy Zenobi-Wong

## Abstract

The vascular tree spans length scales from centimeter to micrometer. Engineering multiscale vasculature, in particular from millimeter vessels to micrometer-size capillaries, represents an unmet challenge and may require the convergence of two or more printing modalities. Leveraging the great advances in light-based biofabrication, we herein introduce a hybrid strategy to tackle this challenge. By combining volumetric printing (VP) and high-resolution two-photon ablation (2PA), we demonstrate the possibility to create complex multiscale organotypic perfusable models with features ranging from mesoscale (VP) to microscale (2PA). To successfully combine these two methods, we first eliminated micrometer-size defects generated during VP process. Due to optical modulation instability of the laser source and self-focusing phenomenon that occurs when the light triggers the photoresin crosslinking, VP printed constructs feature micrometer-size filaments and channels. By optical tuning the refractive index of the photoresin, we demonstrate defect-free VP that can then be combined with 2PA. To facilitate the 2PA process and meet VP requirements, we introduce a purely protein-based photoclick photoresin combining gelatin-norbornene and gelatin-thiol. By optimizing defect-free VP and 2PA processes, we finally demonstrate the possibility to generate complex 3D vasculature-like constructs with features ranging from ~400 μm of VP to ~2 μm of 2PA. This hybrid strategy opens new possibilities to better recapitulate microtissues vasculature and complex architectures, with particular potential for microfluidics and organ/tissue-on-a-chip technologies.

## 1. Introduction

The engineering of organs and tissues on-a-chip holds great promise for biomedical applications such as drug testing and disease modeling.^1–4^ Despite technological advances, current microtissues are much simpler than their native counterparts. In particular, engineering the complex perfusable architectures of the vasculature tree still represents a major challenge in this field. In the past decades, great effort has been made to individually recapitulate the various components of the vasculature tree, from centimeter-scale aorta to micrometer-scale capillaries.^5–6^ However, the field lacks strategies that enable the engineering of multiscale constructs featuring meso- (hundreds of μm) and micro-vasculature (few – tens of μm). Light-based printing offers a broadening and increasingly sophisticated range of techniques for precise fabrication of perfusable tissue architectures.^7^ Volumetric printing (VP) is a novel light-based biofabrication method emerging as a promising technology for such applications, enabling the printing of complex centimeter-size models within seconds. Recent studies have demonstrated the possibility to create hollow, perfusable structures, possibly targeting mesoscale vasculature.^8–10^ However, to fully replicate a multiscale vasculature-like model, VP fall short of reaching microcapillaries size. Another light-based method named two-photon ablation (2PA) offers instead complementary capabilities, being limited in printing time and construct size, but reaching the highest resolution of any biofabrication method (≤ 1 μm). 2PA is based on multiphoton ionization induced by high-intensity pulsed lasers,^11–12^ and has been explored to form cell-instructive microchannels.^13–18^

Seeking to reproduce a multiscale vasculature-like construct, we show for the first time a hybrid VP and 2PA printing technology. In order to successfully combine these two techniques, we first developed a strategy to remove the VP-generated microdefects. As recently shown by Liu et al.,^19^ optical modulation instability (OMI) results in the formation of hydrogel microfilaments and microchannels (void spaces between microfilaments) in the range of 2 −30 μm propagating via self-focusing waveguides (Figure 1A). Therefore, although commonly described as defect-free due to the layer-less printing modality, VP printed constructs have in microfilaments and microchannels a major source of defects which can limit their applications. In fact, while on one hand microchannels can improve diffusion of nutrients into the printed construct, they can also act as physical guidance cues for cells to spread, migrate, align and deposit extracellular matrix (ECM).^19^ Although optimal for anisotropic tissues such as muscle and tendon, this unidirectional microarchitecture is not desirable for all applications requiring an isotropic cell spreading without preferential outgrowth direction or cell confinement in a defined region (i.e., channel wall). In particular for the combination with 2PA, the presence of microchannels hinders high-resolution printing which requires a homogeneous, defect-free material substrate to fully guarantee precise control over hollow architectures.

**Figure 1.**
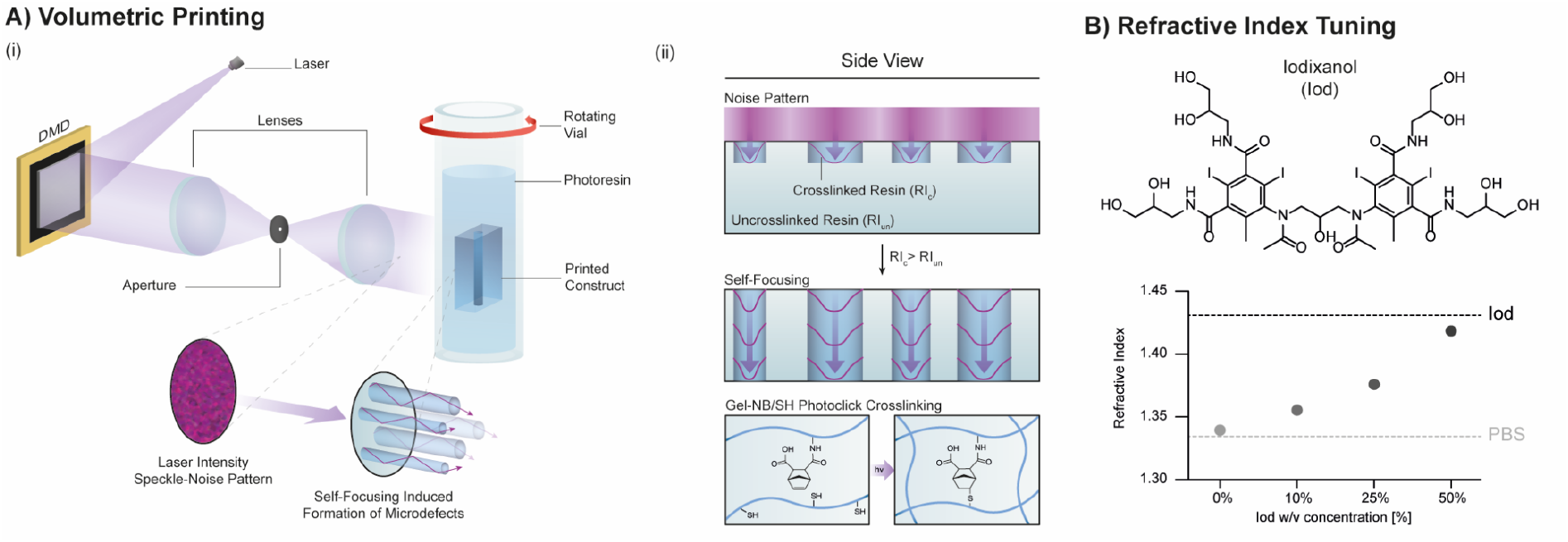
A) (i) Illustration of volumetric printing (VP) method with close-up showing microfilament and microchannel (microdefect) formation due to self-focusing process. (ii) Side view of microdefects formation with details of refractive indices (RI) and photoclick crosslinking mechanism of the gelatin-norbornene (Gel-NB) / gelatin-thiol (Gel-SH) based photoresin. B) Chemical structure of Iodixanol (Iod, top) and linear increase of RI for 2.5% Gel-NB/Gel-SH photoresin containing various Iod %.

Recently, Rackson et al. proposed a strategy to reduce the filamentation effect by reducing the light exposure of the printing process, followed by LED-based uniform exposure (flood illumination) to develop the 3D constructs.^20^ Although theoretically simple and inexpensive, this method requires significant fine-tuning of the exposure time that yields both high resolution and construct stability. In our experience, elimination of microchannels and microfilaments with hydrogel forming photoresins could not be successfully and consistently achieved using this method. In this study, we therefore first introduce an alternative strategy to remove the microdefects by optical tuning of the photoresins, eventually allowing the formation of homogeneous constructs suitable for complex VP / 2PA hybrid printing targeting multiscale organotypic perfusable models.

## 2. Results and Discussion

### 2.1. Refractive Index: How to Limit Self-Focusing Effect

In VP, a laser beam is directed towards a digital micromirror device (DMD) which projects a series of light patterns onto a rotating vial containing the photocrosslinkable material (photoresin) (Figure 1Ai). The projections result in a 3D light-dose accumulation in the photoresin. When the light-dose exceeds the material’s critical gelation threshold, the desired 3D model is formed and can be retrieved with the removal of the uncrosslinked photoresin. However, when reaching the photoresin, the laser beam featuring a speckle-pattern intensity noise causes the formation of microfilaments and microchannels,^19^ herein also generally described as microdefects. This phenomenon originates from the non-linear nature of the photosensitive material which shows a change in refractive index (RI) between its uncrosslinked to crosslinked state. When the local intensity noise maxima crosslink the photoresin, the increase in RI of the crosslinked part results in a self-focusing effect which eventually acts as an optical trap reinforcing the generation of the microfilaments (Figure 1Aii).^21–23^ As the laser noise pattern features low intensity regions between local maxima, uncrosslinked volumes (microchannels) are also formed. Aiming to obtain homogenous, defect-free VP prints suitable for further processing by high-resolution 2PA, we investigated the impact of photoresin RI in limiting the self-focusing effect.

First, to match the requirements of both VP and 2PA, we chose to use a photoresin based on gelatin-norbornene (Gel-NB) and gelatin-thiol (Gel-SH) (Figure 1Aii). As previously shown, thiol-norbornene click-chemistry and gelatin thermal gelation make such photoresin an ideal candidate for VP.^8^ Moreover, as 2PA is known to be more efficient with protein-based systems, ^11^here we introduce for the first time in VP a purely photoclick gelatin-based formulation featuring Gel-SH (Figure 1Aii). For all the experiments, a photoresin composed of 2.5% Gel-NB/SH (with 1:1 NB:SH molar ratio) and 0.05% lithium phenyl-2,4,6-trimethylbenzoylphosphinate (LAP) as photoinitiator (PI) was used.

This initial formulation (RI: 1.3390) was then optically tuned with the addition of iodixanol (Iod) at 10%, 25% or 50%. Iod is a non-toxic, water soluble, non-ionic radiocontrast agent that had also been recently used as RI-matching compound in VP (Figure 1B).^10^ The Iod stock solution (60%) has a significantly higher RI (1.4310) compared to the photoresin and its addition resulted in a linear increase in the photoresins RI up to 1.4195 for Iod 50% (Figure 1B). As described by Kip et al.,^24^ the self-focusing effect due to OMI appears, for a given laser coherence length and intensity, only if the non-linearity (change in RI due to crosslinking) exceeds a certain threshold. We therefore hypothesized that, by adding a compound that increases the overall RI without participating to the crosslinking reaction, we could reduce the change in RI, thus reaching such threshold and eventually hindering microdefects formation (Figure S1, Supporting Information).

### 2.2. Towards Defect-Free Volumetric Printing (VP)

Using the same printing parameters (see Materials and Methods), photoresins containing 0%, 10%, 25% and 50% Iod were used to print a perfusable model (Figure 2Ai). The constructs could be successfully printed and perfused in all photoresin conditions and showed a clear difference in their surface roughness as seen with phase-contrast imaging (Figure 2Ai, close-ups). The microdefects appeared to be reduced with increasing Iod concentration. At 50% Iod, the gel appears macroscopically smooth and homogenous as corroborated by magnified phasecontrast images. To characterize these microdefects in the various formulations, we used confocal reflection imaging (Figure 2Aii). By collecting the light back-scattered from the sample, we could clearly distinguish between void spaces (microchannels, black areas) and gel (microfilaments, cyan areas). Images of the bulk gel and gel-to-PBS solution interface showed a noticeable decrease in the size of the microdefects ultimately resulting in homogeneous, defect-free gel for the photoresin containing 50% Iod. As initially hypothesized, by introducing a high-enough RI homogenous ground level, the system non-linearity (difference in RI between crosslinked and uncrosslinked state) was insufficient to reach the threshold required to induce the self-focusing effect. Quantification of the microdefects size also showed significant reduction with Iod addition from the ~20 μm and ~10 μm for 0% Iod to the 10 μm and ~ 6 μm for 25% Iod microfilaments and microchannels, respectively (Figure 2B). The reduction in microdefects size for intermediate conditions (10% and 25% Iod) could be explained by a distinct phenomenon. The self-focusing effect needs, to occur, an intensity higher than a critical value which is inversely proportional to the RI of the material (Figure S1, Supplementary Information). An increase in the photoresin RI could be therefore expected to induce a denser filamentation as also lower intensity peaks of the laser projection speckle noise pattern could induce self-focusing. To further illustrate the nature of channels and filaments, an infiltration test was conducted using a solution of fluorescently labeled dextran (TRITC-Dex, red). Without Iod, TRITC-Dex solution could easily percolate into the construct via the aligned microchannels (Figure 2C). In contrast, the absence of such void spaces limits solution infiltration to simple diffusion through the hydrogel mesh.

**Figure 2.**
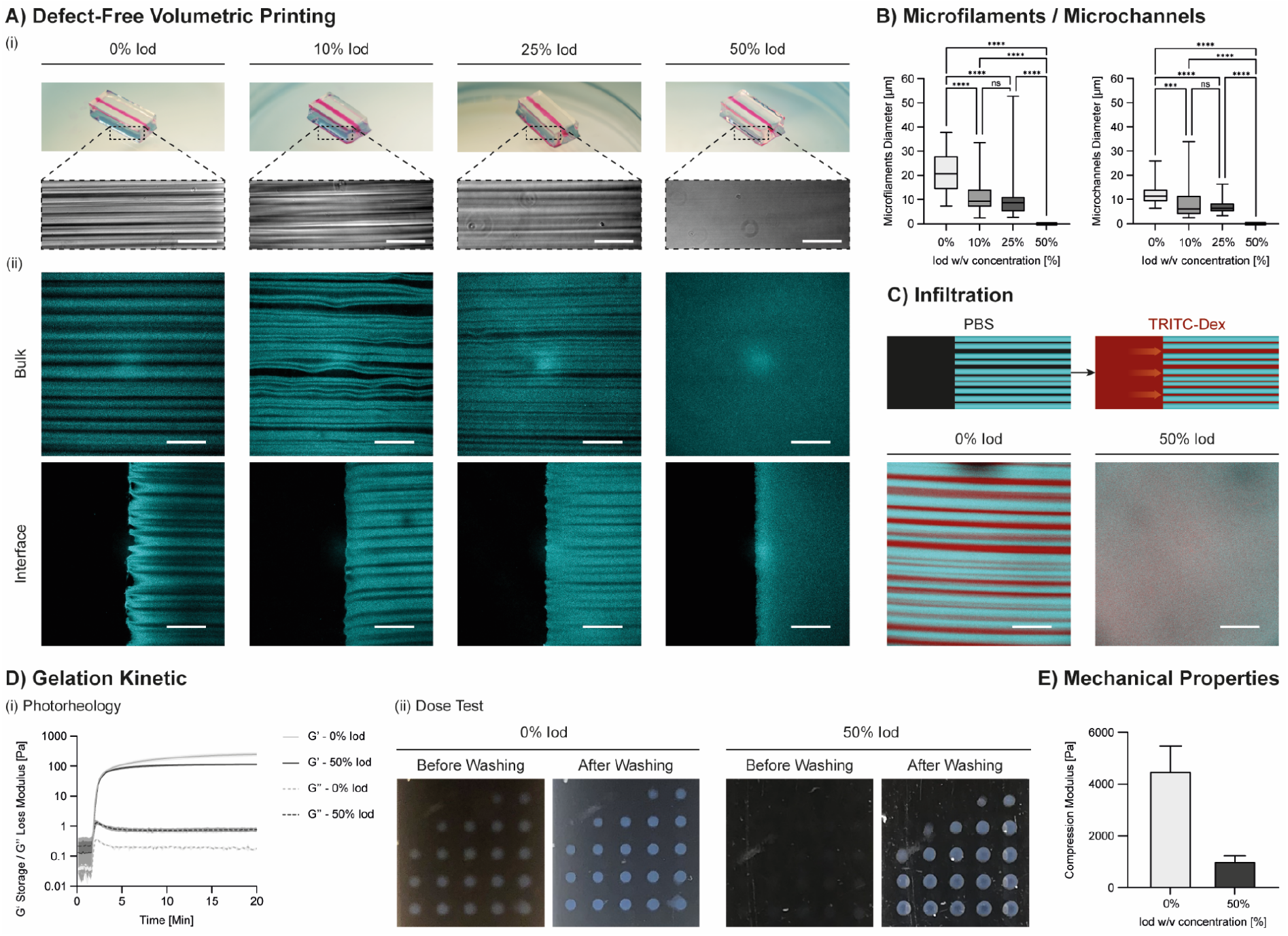
Removal of VP-associated microdefects. A) (i) Volumetric printing of models using different Iod concentrations. Channels were perfused with TRITC-Dex (red). Close up showing phase contrast images of the microfilaments/microchannels. Scale bars: 150 μm (ii) Reflection imaging of the printed constructs taken at the center of the gels (bulk) and at their borders (interfaces) showing a reduction in microfilaments (cyan) and microchannels (black) size with increasing Iod concentration. At 50% Iod, microdefects appeared to be removed. B) Quantitative analysis of microfilaments and microchannels diameters with various Iod concentrations showing size reduction with increased Iod concentration. For 50% Iod, both defects were given a zero value as filamentous structures were discernable. C) Infiltration test with fluorescently labeled TRITC-Dex (40 kDa, red) confirming the presence of perfusable microchannels in 0% Iod condition, and their absence in 50% Iod condition. D) Impact of 50% Iod in gelation kinetic. (i) Photorheological comparison of 50% vs 0% Iod photoresin formulations showed that the presence of Iod determined a reduction in final storage modulus (247 ± 26 Pa for 0% Iod vs 114 ± 7 Pa for 50% Iod) and an increase in loss modulus (0.180 ± 0.008 Pa for 0% Iod vs 0.765 ± 0.085 Pa for 50% Iod), suggesting the formation of a less densely crosslinked network. However, gelation onset appeared to be comparable. (ii) Dose Test confirming comparable critical gelation thresholds (~28.4 mJ cm^3-1^) for the two conditions. Notably, due to negligible difference in RI, crosslinked dots in 50% Iod photoresin could not be visualized without washing out Iod excess. E) Uniaxial compression test on VP printed gels confirmed relevant impact in the mechanical properties of the printed constructs with Iod addition (4494 ± 800 Pa for 0% Iod vs 1014 ± 179 Pa for 50% Iod).

To obtain the desired defect-free printing, the addition of a significant amount of Iod (50%) was proven to be necessary. However, as anticipated, this did not seem to impact the gelation point, thus keeping the printing parameters constant between various formulations. Photorheology measurements conducted at 37°C, showed an essentially identical gelation onset for 0% Iod and 50% Iod formulations (Figure 2Di). This observation was further confirmed with a Dose Test (light dose screening) where the same critical gelation threshold was identified for two photoresins (Figure 2Dii, Figure S2, Supplementary Information). Notably, for the 0% Iod, the crosslinked dots appeared clearly visible before washing out the uncrosslinked resin due to the difference in RI between crosslinked (RI: 1.3445) and uncrosslinked (RI: 1.3390) resin (cause of self-focusing effect and microdefects). In contrast, for the 50% Iod formulation the crosslinked dots were substantially indistinguishable from the uncrosslinked material (unvaried RI: 1.4195) and could be revealed only with washing steps using PBS solution (RI: 1.3345).

Although Iod did not affect the gelation point, the photorheology measurement highlighted a drastic decrease in mechanical properties. It is likely that the large solvation volume of highly concentrated Iod hindered crosslinking completion for 50% Iod photoresin, hence resulting in a lower storage modulus (lower crosslinking density) and increased loss modulus (higher fraction of low crosslinked polymer chains). The mechanical properties of the constructs printed with VP were examined with uniaxial compression testing, confirming a substantial decrease in elastic (compression) modulus with Iod addition (Figure 2E).

### 2.3. Optical Setup: Impact on Microdefects and Resolution

As a light-based method, VP can benefit from a number of optical technologies to further improve or tune the printing process, from lenses setups and DMD to different wavelengths and light sources. In the context of this work, were we aim to generate a multiscale perfusable hydrogel for on-a-chip technologies, VP printing resolution plays a key role. The manipulation of the lens setup is a simple method to improve the printing resolution at the expenses of construct size. In this study, we profited from an open-format VP printer (Readily 3D SA) with interchangeable optics to investigate the impact of an optical setup that doubles the resolution, and consequently decrease by a factor of 2 the projection size. For the sake of clarity, the optical setup used up to this point was termed ‘2x’, whereas ‘1x’ refers to the new optical setup with increased resolution and reduced projection size (Figure 3A). Photoresins with 0% and 50% Iod were tested and confirmed the presence of microdefects in the absence of Iod and homogeneous defect-free constructs with 50% Iod (Figure 3B). Interestingly, we also observed a reduction of the microdefects size by a factor of 2 when using the 1x setup (Figure 3C). This observation is in accordance with the 2-fold reduction in the projection size for the 1x setup. The negative resolution attainable with these optical setups and photoresins were tested by printing hollow conical structures, which were then perfused with high-molecular weight fluorescently labeled dextran (0.5 MDa, FITC-Dex, green) (Figure 3D). The resolution improved from ~420 μm (2x) to ~240 μm (1x) for 0% Iod, and from ~500 μm (2x) to ~380 μm (1x) for 50% Iod. The difference between the two formulations, particularly evident for the 1x, can find a possible explanation in the softness of the matrix formed in the presence of Iod. While for 0% Iod the printed gel is sufficiently stiff to maintain the small diameter cone open, in the 50% Iod the structure tends to collapse as can also be inferred by the thin perfusable tip of the hollow cone.

**Figure 3.**
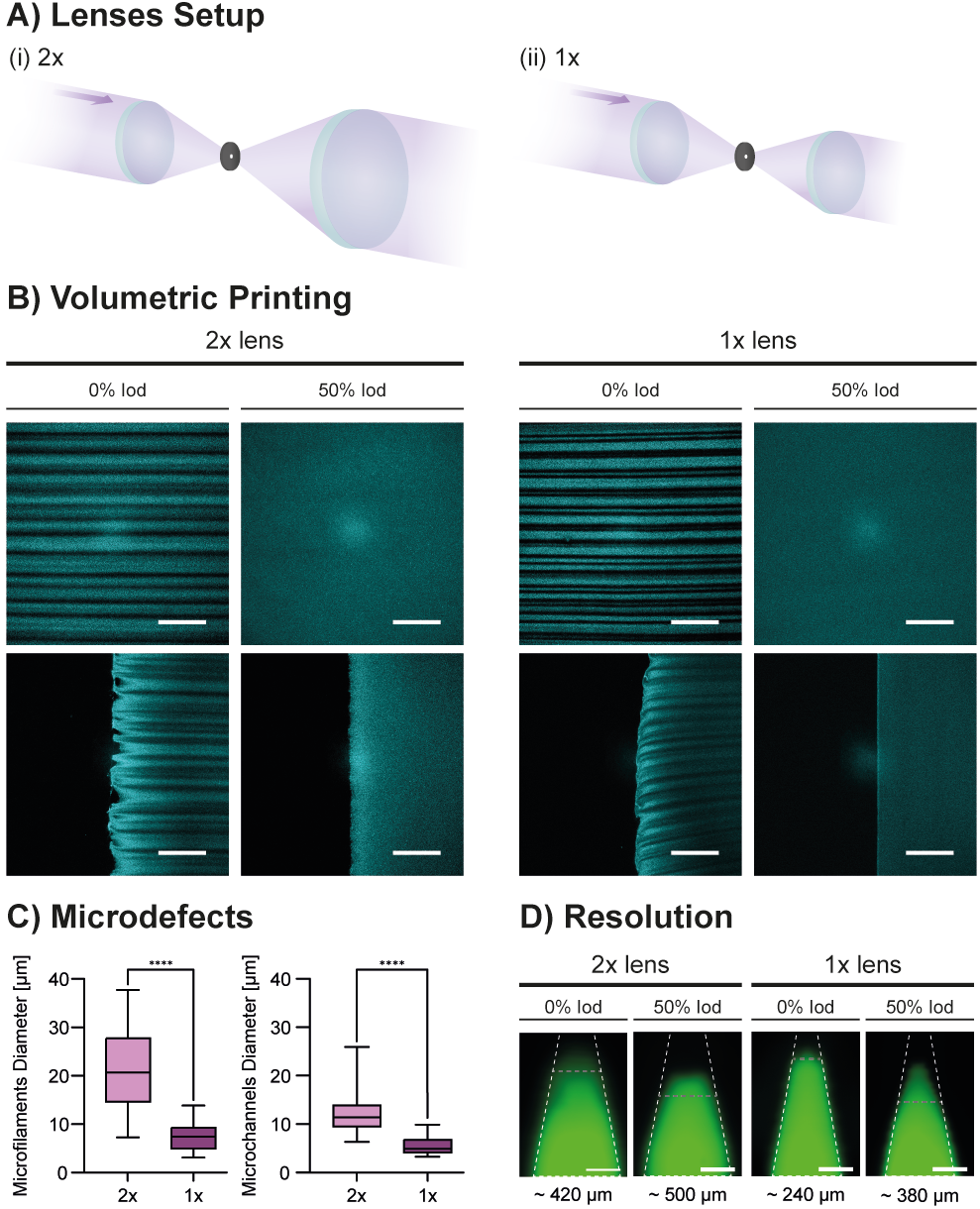
Comparison of VP optical setups. A) Illustration of lenses setups with (i) 2x and (ii) 1x differing in the maximum resolution (double for 1x) and construct/projection size (double for 2x). B) Effect on microdefects of VP printed gels with different lenses setups. Scale bars: 100 μm. C) Quantification of microdefects showing 2-fold reduction in size when using 1x lenses setup compared to 2x. D) Negative resolution of 0% and 50% Iod photoresins performed with VP printing of hollow cones perfused with FITC-Dex (green) showing significant improvement with the use of 1x setup. The maximum resolution was identified with the last point at which the FITC-Dex signal matched the boundaries (grey dashed line) of the theoretical cone dimensions (white dashed line). For 50% Iod – 1x, the FITC-Dex signal was also observed closer to the cone tip, but as the resulting profile did not follow the one of the expected cone (grey dash line), the resolution value was not taken at that point. Scale bars: 300 μm.

### 2.4. Hybrid Printing: Combining Volumetric Printing (VP) with Two-Photon Ablation (2PA)

Having achieved defect-free VP with ~ 400 μm resolution, we then explored the possibility of combining it with 2PA (Figure 4A), a high-resolution method that can enable the fabrication of capillary-sized features. The capabilities of the 2PA setup (see Materials and Methods) were investigated on VP printed gels to identify the optimal parameter working space (printing time, maximum height and resolution). In order to define the areas to scan the laser, regions-of-interest (ROIs) of desired shape were drawn using built-in microscope software functions. As for the study of the microdefects, we used reflection imaging to identify void spaces (ablated areas, black) and printed construct (yellow).

**Figure 4.**
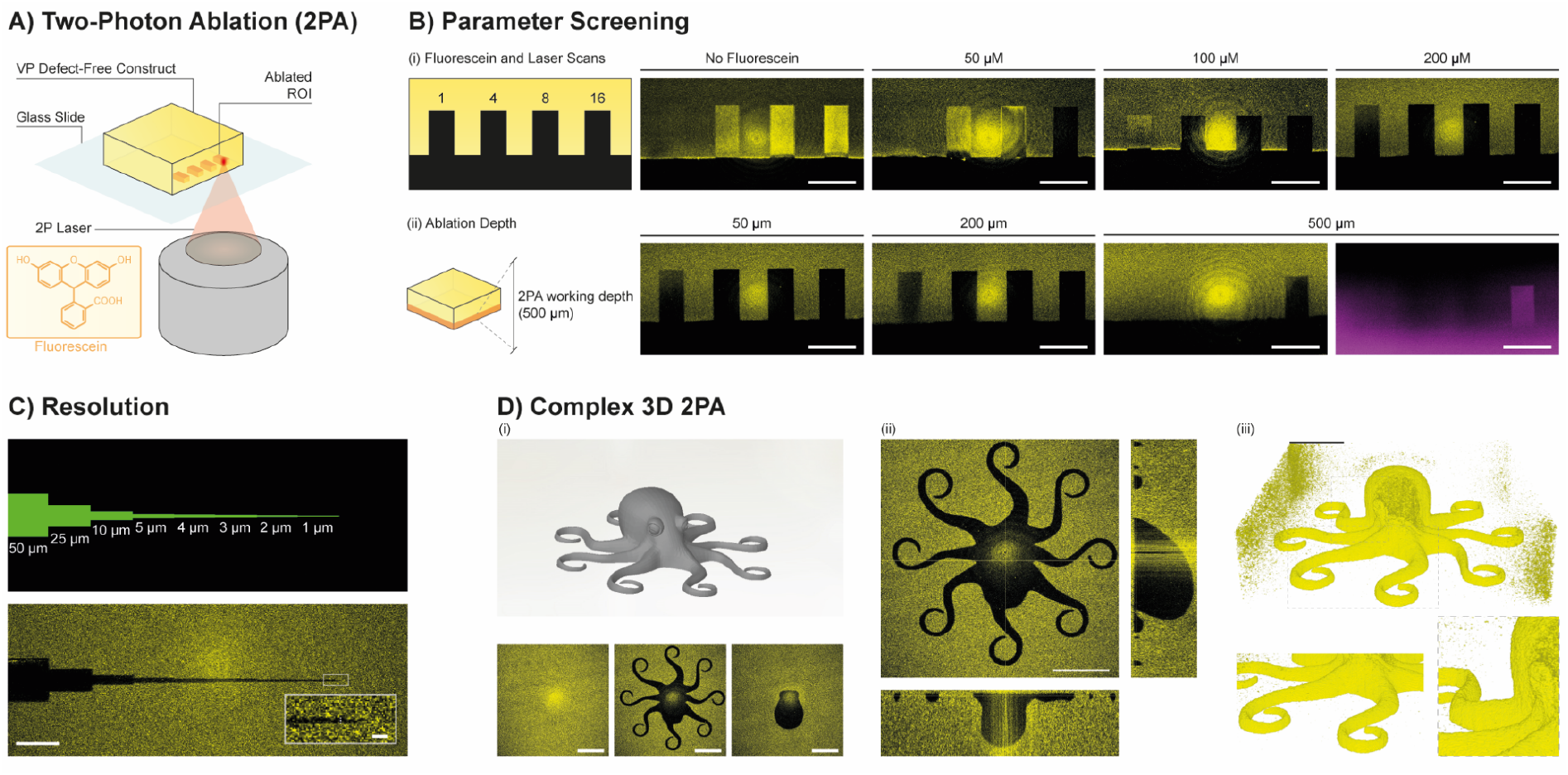
Two-photon ablation (2PA). A) Illustration of 2PA procedure on VP printed gel (yellow). On the bottom-left, chemical structure of fluorescein, used in this work as 2PA-sensitizer. B) (i) Screening of 2PA performance on VP printed gel by varying fluorescein concentration (0, 50 μM, 100 μM and 200 μM) and laser scans (1, 4, 8, 16). Successful ablation of ROIs (in black) was verified using imaging reflection mode. (ii) The best condition (200 μM fluorescein, 16 scans) was used to investigate the maximum 2PA depth (50 μm, 200 μm and 500 μm). With optimized conditions, we found 2PA on VP printed gels to be efficient up to the 500 μm, as also confirmed by the infiltration of TRITC-Dextran (magenta) within the defined ROI. Scale bars: 100 μm. C) 2PA resolution was tested using optimized conditions and a ROI with step-wise decrease in size (green shape, top) and proved to be ~1.8 μm (bottom). Scale bar: 50 μm, close-up 5 μm. D) (i) Model of an octopus (grey, top-left) ablated to show the capability of processing complex 3D models. Images in reflection mode at various heights confirmed accurate spatial ablation. (ii) Orthogonal views of the ablated model showing successful ablation of small details (tentacles) and throughout the whole height. Scale bars: 100 μm. (iii) 3D reconstruction of the ablated octopus with close ups of tentacles and eye details. The artifact of laser reflection at the center of the field-of-view resulted in a distortion on the octopus head during 3D rendering, which is however clearly ablated as can be seen in (ii). Scale bar: 100 μm.

First, we studied the ablation efficiency using different laser scans (number of times the laser scans the same line) and fluorescein concentration at 50 μm depth in the VP construct (Figure 4Bi). Due to its two-photon absorption at the laser peak intensity (205 mW cm^2 −1^ at 780 nm), fluorescein facilitates energy deposition in the matrix acting as a 2PA sensitizer.^25–26^ In addition, fluorescein was chosen as it is a biocompatible, non-expensive and off-the-shelf compound. Fluorescein was found to be fundamental in making the 2PA process effective within the Gel-NB/SH volumetric printed gel. In particular, incubation of the gel in 200 μM fluorescein showed the best results, with efficient ablation even with single laser scanning.

Using a 25x water immersion objective we then used the optimal fluorescein concentration of 200 μM to identify the maximum depth at which we could perform 2PA. We reported successful ablation up to a depth of 500 μm with the use of 16 laser scans (Figure 4Bii). To further confirm proper ablation, we incubated the gel in a solution of TRITC-Dex (magenta) which was shown to infiltrate the void space created during the 2PA process.

Resolution of the 2PA process under optimized conditions was then tested using an ROI with decreasing size, from 50 μm to 1 μm (Figure 4C). The smallest ablated feature was found to be about ~1.8 μm. With the theoretical limit of the optical setup being ~1 μm, the maximum 2PA resolution was almost two times lower likely owed to heat transfer following the high number of scans.

Finally, we ablated an octopus model as a proof-of-concept of our capability to perform complex 3D 2PA (Figure 4D). For this purpose we started from the .stl model of the octopus and converted it into a sequence of layers designed as ROIs using a script previously developed in the lab.^27^ Using the optimized 2PA conditions (200 μM fluorescein, 16 scans) the octopus model was shown to be successfully ablated in all its height and finer structures (Figure 4Di,ii,iii).

### 2.5. Printing of Complex Organotypic Multiscale Models

In the final set of experiments, defect-free VP and optimized high-resolution 2PA were used to generate complex multiscale perfusable models. As 2PA (with the optical setting used in this work) was limited to a maximum working depth of 500 μm, VP printed parts were designed with hollow channels running at this distance from the bottom of the gel (Figure 5A). Starting from the middle of the construct height to facilitate later connection to syringe needles, two parallel channels of 400 μm in diameter were designed to curve downwards reaching the 2PA working volume. The space between the channels was set to 400 μm, approximately corresponding to the field of view of the 2PA optical setup.

**Figure 5.**
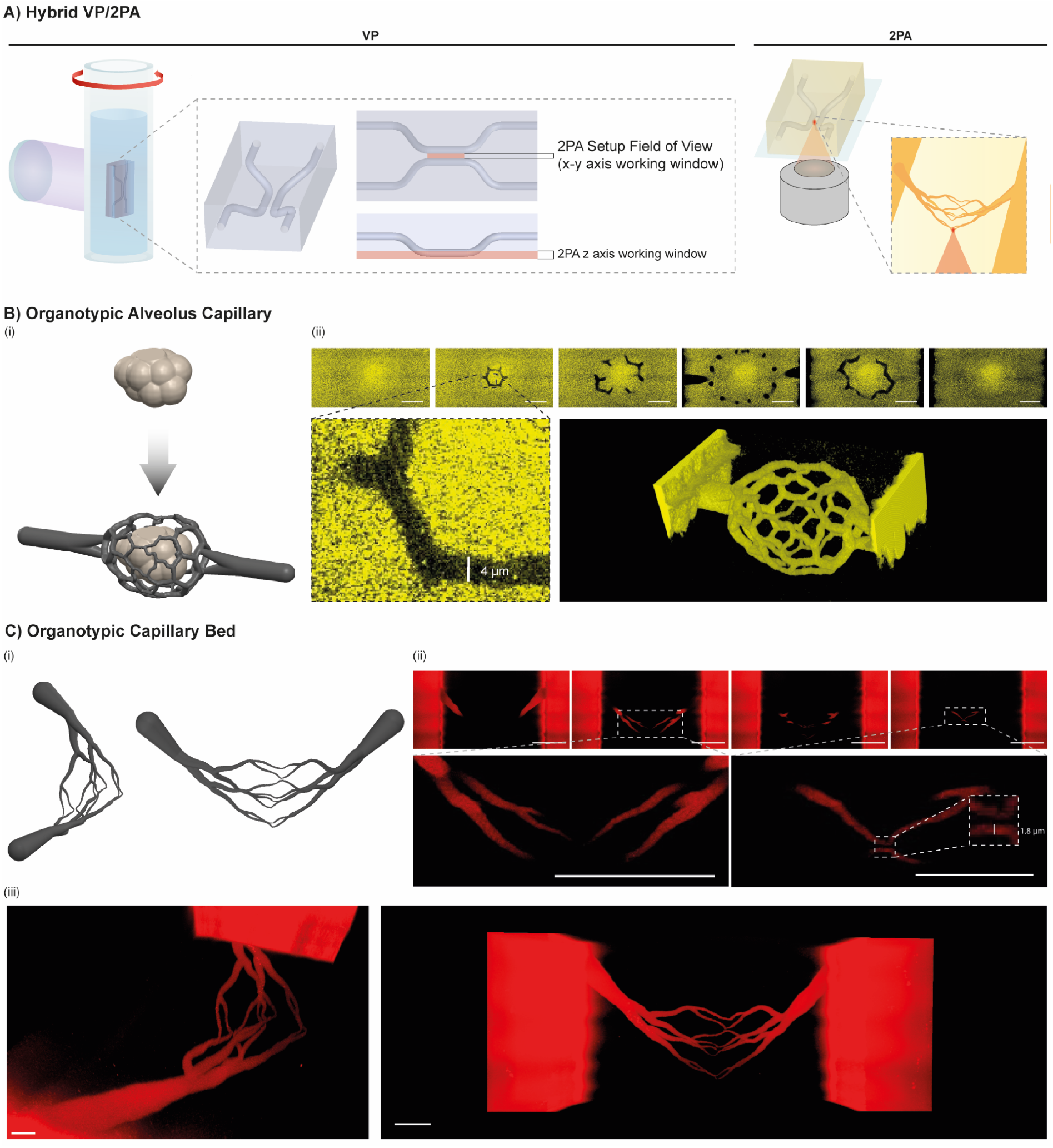
A) Illustration of the hybrid VP/2PA method showing details of the perfusable model printed with VP (left) and following 2PA of capillary-like structures (right). B) Generation (i) and 2PA of complex organotypic capillary network modeled around an alveolus (ii). Successful ablation of the capillary network is shown with reflection images taken at increasing depth (from left to right, scale bar: 50 μm). Close-up showing minimum feature size of ~ 4 μm. On the bottom right a 3D rendering of the ablated model with inverted colors (in yellow the hollow ablated parts). C) (i) Organotypic capillary bed model. (ii) Confocal images at various depth in the gel showing perfusion of gelatin-rhodamine (Gel-Rho) solution. Close up showing details of the fine structures and minimum feature size (~ 1.8 μm). Scale bars: 100μm. (iii) 3D rendering of the perfused capillary bed model connecting mesoscale VP printed channels. Scale bars: 20 μm (left) and 40 μm (right).

Figure 5B shows a first example of a complex multiscale perfusable model produced with the hybrid VP-2PA method. We exploited Hyperganic software platform to generate an organotypic 3D capillary network surrounding an alveolus model of physiological size (~100 μm diameter) (Figure 5Bi). This 3D model was converted into a ROI sequence and ablated between the volumetrically printed parallel channels. Successful combination of defect-free VP and 2PA is shown using reflection imaging at various depths in the gel and 3D reconstruction (Figure 5Bii, Video S1 and S2, Supplementary Information).

To further exploit the potential of this hybrid technique, we ablated a complex organotypic capillary bed featuring branching microcapillaries (Figure 5C). Fluorescently labeled polymer (gelatin-rhodamine) was found to fully percolate into the channels obtained with the 2PA process, thus connecting mesoscale structures obtained with VP with microscale capillary-like structures (Figure 5Cii and iii, Video S3 and S4 Supplementary Information). Notably, the minimum capillary caliper was found to match the resolution limit of this method (~1.8 μm), thus making this example one of the finest capillary-like structure reported so far. The precise fabrication of such highly complex, multiscale models with physiologically relevant native-tissue dimensions opens up new avenues for on-a-chip biomedical applications, such as drug screening and disease modelling.

## 3. Conclusion

In summary, we have introduced a strategy to obtain defect-free VP and leverage it to develop the first VP-based hybrid printing method featuring 2PA. We demonstrated that this method can target complex, multiscale organotypic models, thus opening new avenues for the next generation of on-a-chip technologies. We foresee that the findings of this work will stimulate further research into the optical tuning of the photoresins (i.e., use of various RI matching compounds) and optical setups (i.e., high-resolution VP, 2PA with larger working distance) as well as encourage biological studies in more physiologically relevant tissue models.

## 4. Experimental Section/Methods

### Gelatin-Norbornene (Gel-NB) synthesis

The synthesis protocol was adapted from Rizzo et al..^8, 28^ In short, 25 g of gelatin type A from porcine skin was first dissolved in 0.5 M carbonate–bicarbonate pH 9 buffer at 10% at 40 °C. When completely dissolved, 0.5 g of cis-5-norbornene-endo-2,3-dicarboxylic anhydride (CA) was added to the reaction mixture under vigorous stirring. Every 10 min, 0.5 g of CA was added for a total amount of 2 g. Twenty minutes after the last addition, the solution was diluted 2-fold in prewarmed mQ H2O, and the pH was adjusted to 7 with the addition of HCl 1 M. Then, 2.5 g of NaCl was added and the solution was filter-sterilized (0.2 μm) and dialyzed (12.4 kDa cutoff cellulose tubing) for 4-5 days against mQ H2O at 30 °C with frequent water changes before freeze-drying. The degree of substitution (DS) was calculated using ^1^H NMR (Bruker Ultrashield 400 MHz, 1024 scans), as previously reported,^8, 28^ and found to be ~ 0.091 mmol g^-1^.

### Gelatin-Thiol (Gel-SH) synthesis

The synthesis protocol was adapted from Rizzo et al..^28^ In short, 25 g of gelatin type A from porcine skin was first dissolved in 1.25 L of 150 mM MES pH 4 buffer warmed up to 40 °C for a final concentration of 2%. When completely dissolved, 2.38 g (10 mmol) of 3,3’-dithiobis(proprionohydrazide) (DTPHY) was added to the reaction solution under stirring. When completely dissolved, 3.83 g (20 mmol) of 1-ethyl-3-(3’-dimethylaminopropyl)carbodiimide hydrochloride (EDC) was added, and the reaction was left to proceed overnight at 40 °C. Tris(2-carboxyethyl)phosphine (8.6 g, 30 mmol) was then added to the reaction mixture, and reduction left to proceed for 6 hours in a sealed flask under gentle stirring. Then, 2.5 g of NaCl was added and the solution was filter-sterilized (0.2 μm) and dialyzed (12.4 kDa cutoff cellulose tubing) for 4-5 days against mQ H2O balanced to pH 4.5 with diluted HCl at 30 °C. After dyalisis with frequent changes of acidified water, Gel-SH was finally freeze-dried. Gel-SH was stored under inert atmosphere at −20 °C prior to use. The degree of substitution (DS) was determined by ^1^H NMR (Bruker Ultrashield 400 MHz, 1024 scans) as previously reported,^28^ and found to be ~ 0.276 mmol g^-1^.

### Gelatin-Rhodamine (Gel-Rho) synthesis

Gelatin-methacryloyl was synthesized as previously described.^29^ 1.0 g of GelMA was dissolved in 100 mL of 0.1 M sodium bicarbonate (pH 9.0) at 37 °C overnight. 10.0 mg of rhodamine B isothiocyanate was dissolved in 1.0 mL dimethyl sulfoxide (DMSO) and added to the GelMA solution. The reaction was left to proceed at room temperature overnight. The product was purified by dialysis (3.5 kDa cut-off) against mQ H2O at 40 °C for 3 days with frequent water changes, protected from light, and then freeze dried.

### Photoresin Preparation

Gel-NB and Gel-SH were dissolved in PBS at 37°C with 1:1 molar ratio of NB:SH. Iodixanol (OptiPrepTM, STEMCELL Technologies) stock solution (60%) was added to the Gel-NB/SH mixture for a final concentration of 10%, 25% or 50%. Photoinitiator lithium phenyl-2,4,6-trimethylbenzoylphosphinate (LAP) was added from a stock solution of 2.5% in PBS to obtain a final concentration of 0.05%. Photoresins were prepared with a Gel-NB/SH total polymer concentration of 2.5%, 2.75 % or 3%. Before use, photoresins were filtered through 0.2 μm filters to remove debris and scattering particles.

### Refreactive Index

Refractive indices of the different Gel-NB/SH photoresins were measured using an Abbe refractometer (Kern ORT 1RS, KERN & SOHN GmbH). For measurements of hydrogel refractive indeces, a thin film of photoresin was crosslinked on the measuring prism for 10 minutes in a UV-box (405 nm LEDs, 6.6 mW/cm^2^ intensity).

### Photorheology

Photoreology analyses were carried out on an Anton Paar MCR 302e equipped with a 20 mm parallel plate geometry and glass floor. Omnicure Series1000 lamp (Lumen Dynamics) was used in combination with sequential 400–500 nm and narrow 405 nm bandpass filters (Thorlabs). Gel-NB/SH photoresins were prepared as previously described. All procedures were performed in the dark. Oscillatory measurements were performed in triplicate (n = 3) at 37°C using 74 uL of photoresins at 2% shear rate and 1 Hz frequency with 200 μm gap and 10 s measuring point duration. Measurements were left to proceed in the dark for 1 minute before irradiating the sample with 405 nm light at (60%) 50 mW cm^2-1^ intensity. To prevent the sample from drying all tests were performed in the presence of a wet tissue paper in the chamber.

### Dose Test

A Dose Test was conducted similarly to previously described protocol,^8^ using an open-format printer (Readily3D). In short, photoresins were allowed to thermally gel in quartz cuvettes (CV10Q1400FS, Thorlabs) for 20 min at 4°C prior irradiation. A software built-in function was used to project a 5×5 matrix of dots (0.75 mm diameter, 0.75 mm gap) within a variable range of exposure time and intensity (see Figure S2, Supporting Information). After exposure, cuvettes were imaged before and after washing with PBS.

### Volumetric Printing (VP)

Volumetric printing was performed on an open-format machine, provided by Readily3D, equipped with swappable optics herein described as 2x (larger printing, lower resolution) and 1x (smaller printing, higher resolution). Photoresins were prepared as previously described and allow to thermally gel in the glass vials for 20 min at 4°C prior printing. Printing was conducted with 7.52 mW cm^2-1^ laser intensity and 275 mJ cm^2-1^ light dose, 78.7 ° s^-1^ rotational speed, 273 Hz projection rate. Right after the printing the vials were warmed up in a heating bath (37°C) and uncrosslinked resin was removed with extensive washing in prewarmed PBS with 0.05% LAP. Gels were post-cured in a UV-box (405 nm LEDs, 6.6 mW/cm^2^ intensity) for 10 minutes.

Negative resolution was tested by printing a hollow cone, as previously reported,^8^ perfused with high molecular weight FITC-dextran (0.5 MDa). Fluorescent images were taken on an EVOS M5000 (ThermoFisher). All 3D .stl models to be printed with VP were generated using SolidWorks (Dassault Systèmes).

### Microfilaments/Microchannels Characterization

The microfilaments and microchannels generated during the volumetric printing procedure were imaged in reflection mode using 488 or 633 nm laser line and highly sensitive hybrid (Hyd) detector with HCX IRAPO 25X/0.95NA water immersion objective on Leica TCS SP8 (Leica) confocal microscope. Each condition was tested in triplicate and size of microfilaments and microchannels was measured manually using Fiji. Infiltration experiment was conducted by submerging printed constructs in a solution of 1% TRITC-Dextran (40 kDa) in PBS for 6 hours prior imaging.

### Two-Photon Ablation (2PA)

2PA was performed on a Leica TCS SP8 (Leica) confocal microscope equipped with a Mai Tai two-photon laser (205 mW cm^2-1^ intensity, Spectra-Physics) tuned at 780 nm using HCX IRAPO 25X/0.95NA water immersion objective, 1 μm z-step, 600 Hz scanning, bi-directional scanning, 512×512 format and zoom factor of 1. To avoid drying during ablation procedure the gels were placed in 2 or 8-well glass chambers (Nunc™ Lab Tek™, ThermoScientific) and submerged in PBS with or without fluorescein. 2PA screening was conducted by submerging the VP printed gels for 6 hours in different PBS solutions containing 0, 50, 100 or 200 μM fluorescein. Rectangular-shaped regions-of-interests (ROIs) were designed using LAS X software functionalities and used to test different number of laser scans (1, 4, 8, and 16) and reachable depth in the gel (50, 200 and 500 μm). Resolution test was performed with optimized conditions (16 scans, 200 μM fluorescein) using a ROI designed to have a step-wise decrease in size, from 50 to 1 μm (see Figure 4C).

Ablation of the octopus 3D model was performed using a script previously developed in our lab (available at https://github.com/nbroguiere/F2P2),^27^ which converts an .stl file into a stack of ROIs executable by the LAS X software in Live Data Mode. Ablated construct was imaged with 633 nm backscattered light (reflection mode), 0.5 μm z-step, and elaborated with Fiji.

Capillary bed and alveoli models were generated using Hyperganic (Hyperganic Group) and ablated using the same procedure described above. After ablation, samples were perfused with 0.5% Gel-Rho solution in PBS and imaged using Leica SP8 confocal microscope. 3D rendering and videos were generated with Fiji or Imaris.

## Acknowledgements

The authors acknowledge ETH ScopeM imaging facility for their assistance.

## Supporting Information

**Figure S1.**
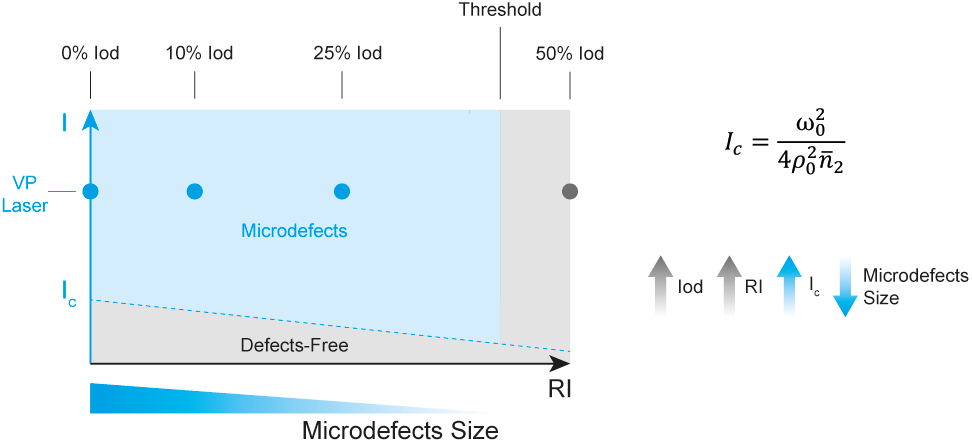
Illustration representing the two phenomena involved in the reduction and disappearance of microdefects. As displayed in the equation,^30^ the critical light intensity (Ic) above which self-focusing effects occurs is inversely proportional to the refreactive index (RI, n2) of the material. Therefore, given the VP laser intensity, with an increase in iodixanol (Iod) concentration – and thus an increase in refractive index (RI) – the self-focusing effect occurs at lower intense speckles determining smaller microdefects. This phenomenon, observable with 10% and 25% Iod (see Figure 2A), occurs till a critical threshold at which the RI is so uniformly high that the RI change between crosslinked and uncrosslinked state is negligible and does not trigger microdefects formation via self-focusing effect. This defect-free region where a defects-free gel is formed was obtained using 50% Iod. The critical intensity I_c_ is also related to other parameters such as laser beam waist (ω) and Raylegh length (τ).

**Figure S2.**
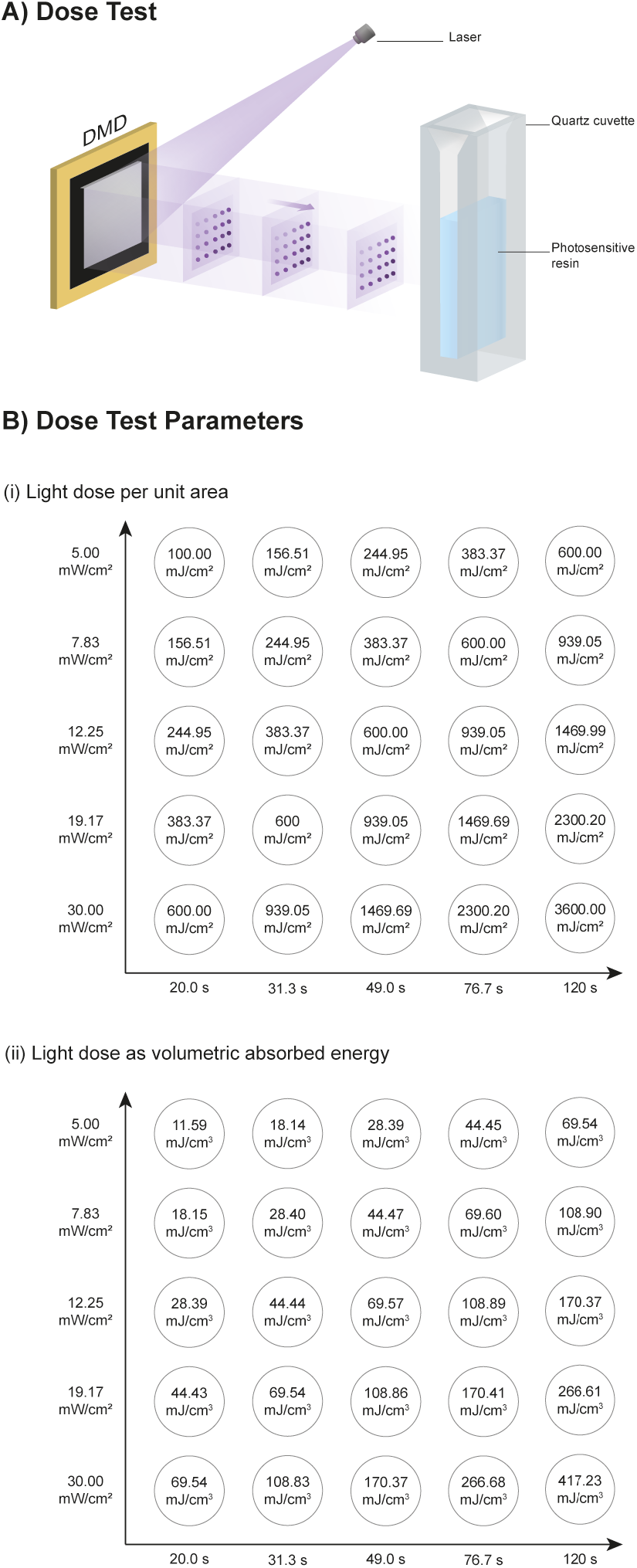
A) Illustration of Dose Test procedure with dots of various intensities projected, for different time, towards a static (non-rotating) quartz cuvette filled with photoresin. B) Parameters used for the matrix of dots with light dose in aereal (i) and volumetric units (ii).

**Video S1**. Reflection imaging z-stack into VP printed gel (yellow) showing ablated regions (black) of the organotypic 3D capillary modeled around an alveolus-like shape.

**Video S2**. Animation of 3D rendered alveolus capillary network (yellow).

**Video S3**. Fluorescent imaging z-stack into VP printed gel (black) showing capillary-bed ablated regions perfused with gelatin-rhodamine solution (red).

**Video S4**. Animation of 3D rendered capillary-bed network (red).

## References

1. Bhatia, S. N.; Ingber, D. E., Microfluidic organs-on-chips. Nature Biotechnology 2014, 32 (8), 760–772.

2. Leung, C. M.; de Haan, P.; Ronaldson-Bouchard, K.; Kim, G.-A.; Ko, J.; Rho, H. S.; Chen, Z.; Habibovic, P.; Jeon, N. L.; Takayama, S.; Shuler, M. L.; Vunjak-Novakovic, G.; Frey, O.; Verpoorte, E.; Toh, Y.-C., A guide to the organ-on-a-chip. Nature Reviews Methods Primers 2022, 2 (1), 33.

3. Qiu, Y.; Ahn, B.; Sakurai, Y.; Hansen, C. E.; Tran, R.; Mimche, P. N.; Mannino, R. G.; Ciciliano, J. C.; Lamb, T. J.; Joiner, C. H.; Ofori-Acquah, S. F.; Lam, W. A., Microvasculature-on-a-chip for the long-term study of endothelial barrier dysfunction and microvascular obstruction in disease. Nature Biomedical Engineering 2018, 2 (6), 453–463.

4. Hsu, Y.-H.; Moya, M. L.; Hughes, C. C. W.; George, S. C.; Lee, A. P., A microfluidic platform for generating large-scale nearly identical human microphysiological vascularized tissue arrays. Lab on a Chip 2013, 13 (15), 2990–2998.

5. Fleischer, S.; Tavakol, D. N.; Vunjak-Novakovic, G., From Arteries to Capillaries: Approaches to Engineering Human Vasculature. Advanced Functional Materials 2020, 30 (37), 1910811.

6. Xie, R.; Zheng, W.; Guan, L.; Ai, Y.; Liang, Q., Engineering of Hydrogel Materials with Perfusable Microchannels for Building Vascularized Tissues. Small 2020, 16 (15), 1902838.

7. Lee, M.; Rizzo, R.; Surman, F.; Zenobi-Wong, M., Guiding Lights: Tissue Bioprinting Using Photoactivated Materials. Chemical Reviews 2020, 120 (19), 10950–11027.

8. Rizzo, R.; Ruetsche, D.; Liu, H.; Zenobi-Wong, M., Optimized Photoclick (Bio)Resins for Fast Volumetric Bioprinting. Advanced Materials 2021, 33 (49), 2102900.

9. Bernal, P. N.; Delrot, P.; Loterie, D.; Li, Y.; Malda, J.; Moser, C.; Levato, R., Volumetric Bioprinting of Complex Living-Tissue Constructs within Seconds. Advanced Materials 2019, 31 (42), 1904209.

10. Bernal, P. N.; Bouwmeester, M.; Madrid-Wolff, J.; Falandt, M.; Florczak, S.; Rodriguez, N. G.; Li, Y.; Größbacher, G.; Samsom, R.-A.; van Wolferen, M.; van der Laan, L. J. W.; Delrot, P.; Loterie, D.; Malda, J.; Moser, C.; Spee, B.; Levato, R., Volumetric Bioprinting of Organoids and Optically Tuned Hydrogels to Build Liver-Like Metabolic Biofactories. Advanced Materials 2022, 34 (15), 2110054.

11. Pradhan, S.; Keller, K. A.; Sperduto, J. L.; Slater, J. H., Fundamentals of Laser-Based Hydrogel Degradation and Applications in Cell and Tissue Engineering. Advanced Healthcare Materials 2017, 6 (24), 1700681.

12. Schaffer, C. B.; Brodeur, A.; Mazur, E., Laser-induced breakdown and damage in bulk transparent materials induced by tightly focused femtosecond laser pulses. Measurement Science and Technology 2001, 12 (11), 1784–1794.

13. Rayner, S. G.; Howard, C. C.; Mandrycky, C. J.; Stamenkovic, S.; Himmelfarb, J.; Shih, A. Y.; Zheng, Y., Multiphoton-Guided Creation of Complex Organ-Specific Microvasculature. Advanced Healthcare Materials 2021, 10 (10), 2100031.

14. Mark, A. S.-S.; Man-Chi, L.; Yuelong, W.; Mehmet Fatih, Y. In Multi-photon microfabrication of three-dimensional capillary-scale vascular networks, Proc.SPIE, 2017; p 101150L.

15. Sarig-Nadir, O.; Livnat N Fau - Zajdman, R.; Zajdman R Fau - Shoham, S.; Shoham S Fau - Seliktar, D.; Seliktar, D., Laser photoablation of guidance microchannels into hydrogels directs cell growth in three dimensions. (1542–0086 (Electronic)).

16. Applegate, M. B.; Coburn, J.; Partlow, B. P.; Moreau, J. E.; Mondia, J. P.; Marelli, B.; Kaplan, D. L.; Omenetto, F. G., Laser-based three-dimensional multiscale micropatterning of biocompatible hydrogels for customized tissue engineering scaffolds. Proceedings of the National Academy of Sciences 2015, 112 (39), 12052–12057.

17. Lee, B. L.-P.; Jeon, H.; Wang, A.; Yan, Z.; Yu, J.; Grigoropoulos, C.; Li, S., Femtosecond laser ablation enhances cell infiltration into three-dimensional electrospun scaffolds. Acta Biomaterialia 2012, 8 (7), 2648–2658.

18. Enrico, A.; Voulgaris, D.; Östmans, R.; Sundaravadivel, N.; Moutaux, L.; Cordier, A.; Niklaus, F.; Herland, A.; Stemme, G., 3D Microvascularized Tissue Models by Laser-Based Cavitation Molding of Collagen. Advanced Materials 2022, 34 (11), 2109823.

19. Liu, H.; Chansoria, P.; Delrot, P.; Angelidakis, E.; Rizzo, R.; Ruetsche, D.; Applegate, L. A.; Loterie, D.; Zenobi-Wong, M., Filamented Light (FLight) Biofabrication of Highly Aligned Tissue - engineered Constructs. Advanced Materials 2022, n/a (n/a), 2204301.

20. Rackson, C. M.; Toombs, J. T.; De Beer, M. P.; Cook, C. C.; Shusteff, M.; Taylor, H. K.; McLeod, R. R., Latent image volumetric additive manufacturing. Opt. Lett. 2022, 47 (5), 1279–1282.

21. Kewitsch, A. S.; Yariv, A., Self-focusing and self-trapping of optical beams upon photopolymerization. Opt. Lett. 1996, 21 (1), 24–26.

22. Burgess, I. B.; Shimmell, W. E.; Saravanamuttu, K., Spontaneous Pattern Formation Due to Modulation Instability of Incoherent White Light in a Photopolymerizable Medium. Journal of the American Chemical Society 2007, 129 (15), 4738–4746.

23. Biria, S.; Malley, P. P. A.; Kahan, T. F.; Hosein, I. D., Tunable Nonlinear Optical Pattern Formation and Microstructure in Cross-Linking Acrylate Systems during Free-Radical Polymerization. The Journal of Physical Chemistry C 2016, 120 (8), 4517–4528.

24. Kip, D.; Soljacic, M.; Segev, M.; Eugenieva, E.; Christodoulides, D. N., Modulation Instability and Pattern Formation in Spatially Incoherent Light Beams. Science 2000, 290 (5491), 495–498.

25. Skylar-Scott, M. A.; Liu, M.-C.; Wu, Y.; Dixit, A.; Yanik, M. F., Guided Homing of Cells in Multi-Photon Microfabricated Bioscaffolds. Advanced Healthcare Materials 2016, 5 (10), 1233–1243.

26. Scott, M. A.; Wissner-Gross, Z. D.; Yanik, M. F., Ultra-rapid laser protein micropatterning: screening for directed polarization of single neurons. Lab on a Chip 2012, 12 (12), 2265–2276.

27. Broguiere, N.; Lüchtefeld, I.; Trachsel, L.; Mazunin, D.; Rizzo, R.; Bode, J. W.; Lutolf, M. P.; Zenobi-Wong, M., Morphogenesis Guided by 3D Patterning of Growth Factors in Biological Matrices. Advanced Materials 2020, 32 (25), 1908299.

28. Rizzo, R.; Bonato, A.; Chansoria, P.; Zenobi-Wong, M., Macroporous Aligned Hydrogel Microstrands for 3D Cell Guidance. ACS Biomaterials Science & Engineering 2022, 8 (9), 3871–3882.

29. Kessel, B.; Lee, M.; Bonato, A.; Tinguely, Y.; Tosoratti, E.; Zenobi-Wong, M., 3D Bioprinting of Macroporous Materials Based on Entangled Hydrogel Microstrands. Advanced Science 2020, 7 (18), 2001419.

30. Boyd, R. W.; Prato, D., Nonlinear Optics. Elsevier Science: 2008.

